# Sleep and mental health independently affect cognitive performance in university students

**DOI:** 10.1101/2025.09.30.679650

**Authors:** Aina E Roenningen, Devan Gill, Brianne A Kent

**Author notes:** Correspondence: Brianne A Kent, Simon Fraser University Department of Psychology 8888 University Drive Burnaby, BC Canada.

## Abstract

**Objectives:** Young adults experience the highest rates of mental health disorders of any age group. Given that mental health disorders are associated with sleep disturbances and cognitive impairments, we investigated whether sleep moderates the effects on cognition.

**Methods:** University students (N=89; aged 18-30 years) remotely monitored their sleep for seven consecutive days using wrist actigraphy and sleep diaries. On day seven, participants completed cognitive testing and mental health questionnaires. Cognitive tests included the Psychomotor Vigilance Task (PVT), Cambridge Neuropsychological Test Automated Battery (CANTAB), and the Mnemonic Similarity Task (MST). CANTAB’s Delayed Matching to Sample (DMS) and MST are designed to tax pattern separation, a computational mechanism supporting encoding of similar experiences as distinct representations. Beck’s Depression Inventory and Beck’s Anxiety Inventory assessed mental health.

**Results:** Eighty participants (mean age: 20.13±2.00) were included in the analyses. Most participants reported mild to severe depressive and anxiety symptoms. Depressive symptoms were correlated with wake-up time (ρ*=.*35, *p=.*002) as well as PVT (ρ*=.*26, *p=.*02) and DMS (ρ*=.*24, *p=*.04) performance. Bedtime was correlated with performance on MST (*r=-.*29, *r=.*02) and DMS (ρ*=.*25, *p=.*03), while wake-up time was correlated with performance on MST (*r=- .*31, *p=.*01) and DMS (ρ*=.*28, *p=.*01). Sleep did not moderate the effects of mental health on cognitive performance.

**Conclusion:** Cognitive tests taxing pattern separation are sensitive to depressive symptoms and sleep timing. While students face a disproportionate burden of mental health disorders compromising cognitive functioning, improving sleep quality may offer a partial, though not moderating, pathway to alleviating these cognitive impairments.

## 1. Introduction

Mental health disorders affect quality of life and increase the risk of illness, disability, and mortality [1,2]. Worldwide, one in eight people experience a mental health disorder, with depression and anxiety being the most prevalent conditions [3]. Depressive disorders include mood-related conditions characterized by persistent emotions of sadness, despair, and disinterest in once enjoyable activities [4]. Among the most prevalent of these disorders is Major Depressive Disorder (MDD), with a lifetime prevalence of 12% [5]. Anxiety disorders encompass mental health conditions characterized by persistent and intense emotions of fear and worry, often occurring alongside physical symptoms including restlessness and impaired concentration [4]. Common anxiety disorders such as Panic Disorder, Social Anxiety Disorder, and Generalized Anxiety Disorder (GAD) often co-occur with depression [6].

Although symptoms of anxiety and depression are complex and are expressed differently across individuals, sleep disorders (e.g., insomnia) and sleep disturbances frequently co-occur with anxiety and depression [7]. Those who are struggling with sleep disorders or who experience disrupted sleep are also more likely to develop depressive and anxiety disorders [8,9], suggesting a bi-directional relationship between sleep and mental health.

In college students, a delayed sleep time is common and a predictor of mental health outcomes, likely due to the resulting circadian misalignment and insufficient sleep duration [10–12]. Mental health disorders (e.g., anxiety and depression) and low quality or quantity of sleep have been linked to poor cognitive performance [10,13–15]. For example, research has shown that anxiety, depression, and sleep deprivation are associated with episodic memory deficits, including impaired performance on tests of pattern separation [16,17,20,21]. Pattern separation is a computational process hypothesized to enable the formation of distinct neural representations from similar inputs [18,19], and is critically important for effective episodic memory. Anxiety, depression, and sleep deprivation may impair pattern separation through suppression of neural plasticity in the hippocampus [22,23], which in rodent models has been shown to be critical for pattern separation [17,24].

As college students frequently experience mental health disorders and disordered sleep, the present research aimed to explore whether sleep moderates the effects of depression and anxiety on cognitive performance. We hypothesized that the low sleep quality and quantity associated with mental health symptoms negatively affects performance on cognitive tests, particularly those designed to assess pattern separation. If sleep is shown to moderate the effect of mental health symptoms on cognition, then interventions to promote sleep could be explored to support college students experiencing cognitive impairments associated with depression or anxiety.

## 2. Methods

### 2.1 Participants

The study received approval from the Research Ethics Board (REB) at Simon Fraser University (SFU; protocol #30000539), adhering to the Tri-Council Policy Statement on Ethical Conduct for Research Involving Humans. Subjects were 89 undergraduate students recruited via the SFU Psychology Research Participation System, and through word-of-mouth and poster advertisements. Participants completed an online consent form and questionnaire, which collected demographic information as well as information about mental, neurological, and physical health. We recruited individuals who were between 18 - 30 years of age, who could understand written and oral English. A summary of participant demographics is presented in Table 1.

**Table 1.**
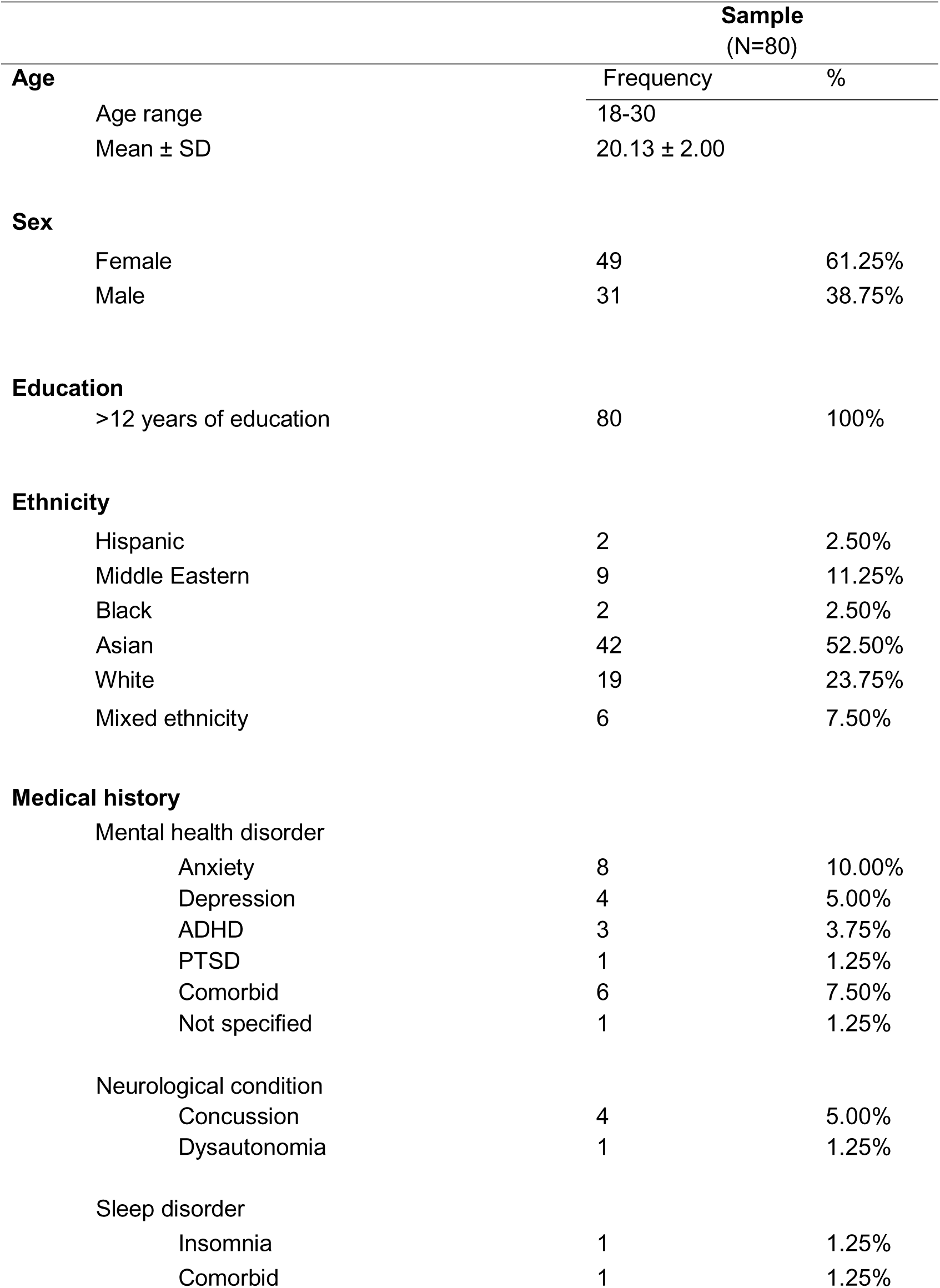
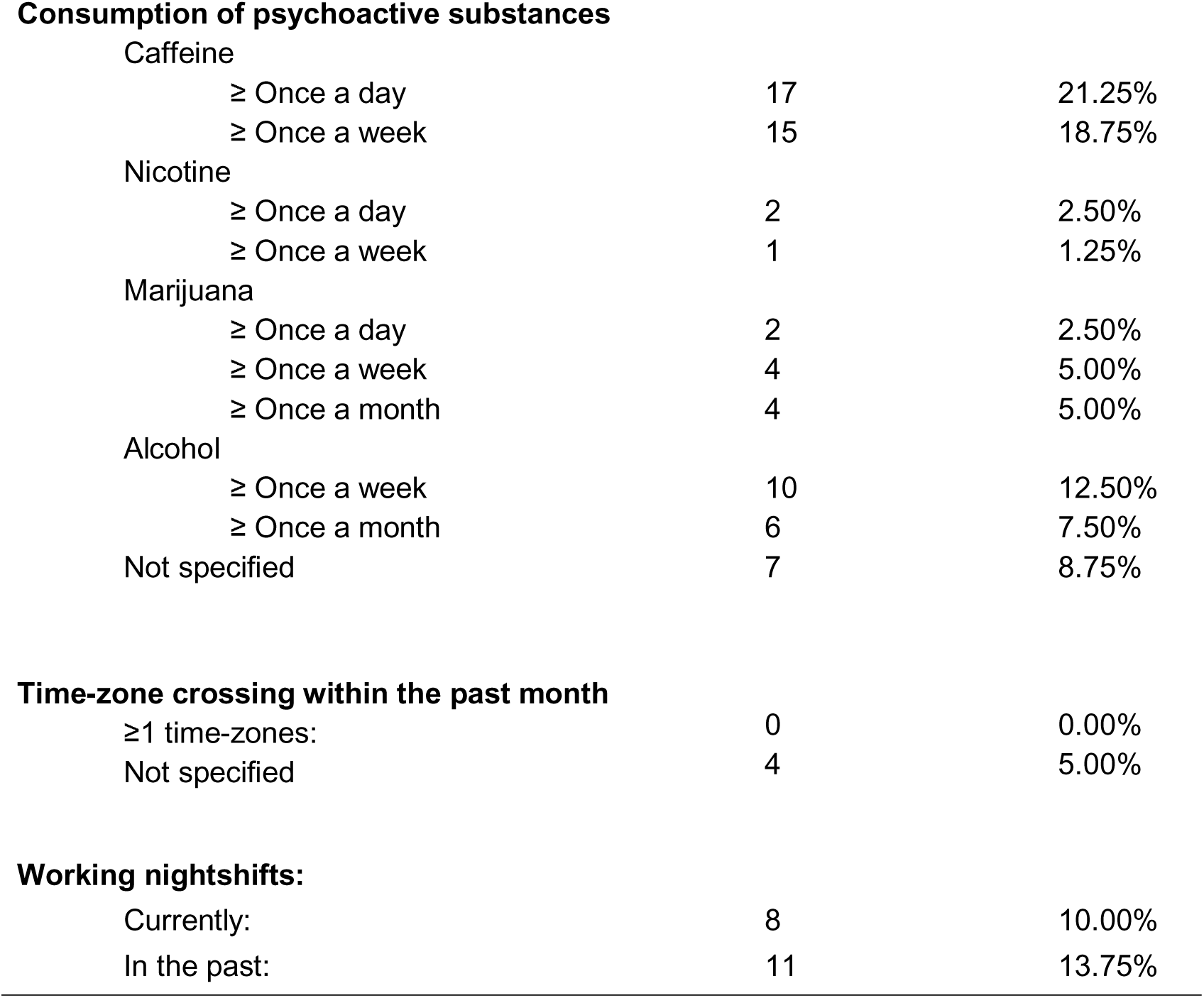
Participant characteristics

|  |  | **Sample**<br>**(N=80)** |  |
| --- | --- | --- | --- |
| **Age** |  | Frequency | % |
| Age range |  | 18-30 |  |
| Mean ± SD | | 20.13 ± 2.00 | |
| **Sex** |  |  |  |
| Female |  | 49 | 61.25% |
| Male |  | 31 | 38.75% |
| **Education** |  |  |  |
| >12 years of education |  | 80 | 100% |
| **Ethnicity** |  |  |  |
| Hispanic |  | 2 | 2.50% |
| Middle Eastern |  | 9 | 11.25% |
| Black |  | 2 | 2.50% |
| Asian |  | 42 | 52.50% |
| White |  | 19 | 23.75% |
| Mixed ethnicity |  | 6 | 7.50% |
| **Medical history** |  |  |  |
| Mental health disorder |  |  |  |
| Anxiety |  | 8 | 10.00% |
| Depression |  | 4 | 5.00% |
| ADHD |  | 3 | 3.75% |
| PTSD |  | 1 | 1.25% |
| Comorbid |  | 6 | 7.50% |
| Not specified |  | 1 | 1.25% |
| Neurological condition |  |  |  |
| Concussion |  | 4 | 5.00% |
| Dysautonomia |  | 1 | 1.25% |
| Sleep disorder |  |  |  |
| Insomnia |  | 1 | 1.25% |
| Comorbid |  | 1 | 1.25% |

**Consumption of psychoactive substances**
|  |  |  |
| --- | --- | --- |
| Caffeine |  |  |
| ≥ Once a day | 17 | 21.25% |
| ≥ Once a week | 15 | 18.75% |
| Nicotine |  |  |
| ≥ Once a day | 2 | 2.50% |
| ≥ Once a week | 1 | 1.25% |
| Marijuana |  |  |
| ≥ Once a day | 2 | 2.50% |
| ≥ Once a week | 4 | 5.00% |
| ≥ Once a month | 4 | 5.00% |
| Alcohol |  |  |
| ≥ Once a week | 10 | 12.50% |
| ≥ Once a month | 6 | 7.50% |
| Not specified | 7 | 8.75% |

**Time-zone crossing within the past month**
|  |  |  |
| --- | --- | --- |
| ≥1 time-zones: | 0 | 0.00% |
| Not specified | 4 | 5.00% |

**Working nightshifts:**
|  |  |  |
| --- | --- | --- |
| Currently: | 8 | 10.00% |
| In the past: | 11 | 13.75% |

### 2.2 Procedure

Participants in the study attended two in-person visits, scheduled 7 days apart between 10:00 am and 3:00 pm. The visits were scheduled during academic semesters and not during the exam period or semester break (September 2021 – February 2023). During the first visit, participants were provided with wrist actigraphy to monitor their rest-activity over the week. On the seventh day, they returned to the lab for the second visit, during which they completed cognitive testing and two mental health questionnaires (Figure 1).

**Figure 1.**
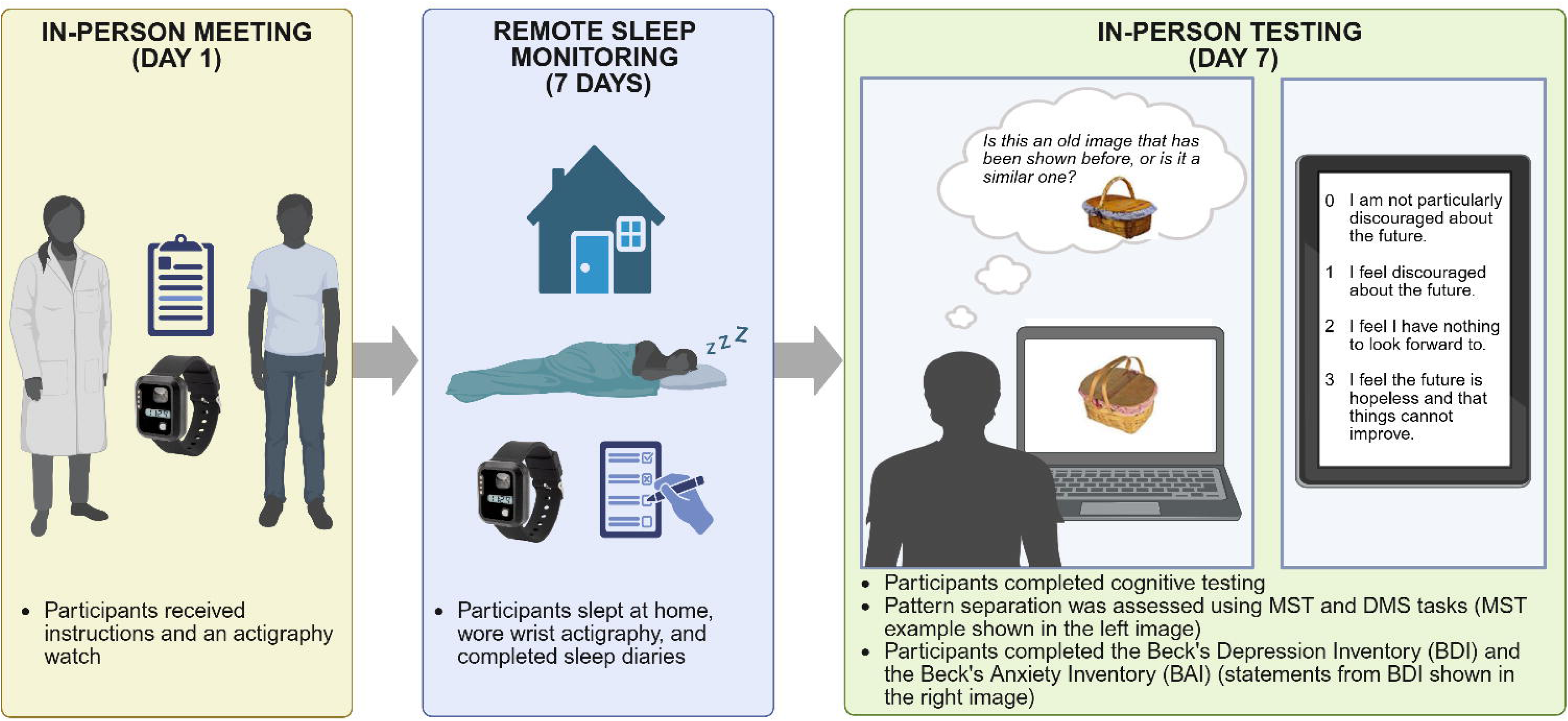
Illustration of the procedure, including two example images used in the MST.^31^ Created with BioRender.com. Abbreviations: MST, Mnemonic Similarity Task; DMS, Delayed Matching to Sample; BDI, Beck’s Depression Inventory; BAI, Beck’s Anxiety Inventory.

### 2.3 Sleep Assessments

Participants wore wrist actigraphy (ActTrust2 model AT20101, Condor Instruments, Sao Paulo, Brazil) on their non-dominant hand and filled out an electronic sleep diary (modified from [25]), over seven consecutive days. They were instructed to use the event button on the wrist actigraphy to record the time they went to bed and the time they woke up.

### 2.4 Cognitive testing

#### 2.4.1 Psychomotor Vigilance Task (PVT)

The PVT is a measure of reaction time and sustained attention [26,27]. We used the 5-min PVT version (Version 1.2.0., NASA, Washington, D.C., USA) [28,29]. Participants completed the task on an Apple iPad (OS 15.4.1, model A2602). In a small rectangular area, a millisecond counter was presented and appeared at random inter-stimulus intervals (ISI), in the range of 2 to 10 s [30]. The millisecond counter rapidly increased from zero in each trial. Participants were instructed to tap their dominant hand’s thumb on the screen as fast as possible once they detected the millisecond counter. Participants completed practice trials before the experimental trials.

#### 2.4.2 Mnemonic Similarity Task (MST)

The MST (version 0.96) is a cognitive task designed to assess pattern separation [31]. We used the computerized task pace, the 3-choice version, and Set 1 stimuli for the MST. The task included two phases. During phase 1 of the task, participants were presented with pictures of everyday objects. They were instructed to indicate via keyboard presses whether the object that appeared belonged indoors or outdoors [31,32]. We did not inform participants about the upcoming memory test in phase 2. Phase 2 had 192 trials and participants were shown pictures of everyday objects. During phase 2, participants were instructed to respond “old,” “similar,” or “new” to indicate whether the objects were a) the exact same as an object seen in phase 1 (i.e., old/target), b) similar to an object seen in phase 1 (i.e., similar/lure), or c) a new image not shown during phase 1 (i.e., new/foil). Of the 192 trials, one-third (64) were targets, one-third (64) were lures, and one-third (64) were foils. The lures included 5 levels of similarity. The similarity levels ranged from L1 (highly similar) to L5 (least similar). In both phases of the task, all pictures were shown with a 2000 ms duration. There was a 500 ms inter-stimulus interval between the presentation of each image.

#### 2.4.3 Cambridge Neuropsychological Test Automated Battery (CANTAB)

Participants completed 5 cognitive tasks from CANTAB, administered in the following order: Delayed Matching to Sample (DMS), Motor Screening Task (MOT), Paired Associates Learning (PAL), Reaction Time (RTI), and Spatial Working Memory (SWM). All tests were conducted using an iPad running OS 15.4.1.

*DMS.* The DMS assessed visual matching and short-term visual recognition memory for abstract and non-verbalizable patterns [33]. A complex sample pattern was shown for 4.5 s, followed by 4 choice patterns that had different structures and colors. Participants were asked to identify the choice pattern that was identical to the sample pattern. The sample pattern and choice patterns appeared simultaneously or there was a delay of 0, 4, or 12 s before the choices were shown. *PAL.* The PAL task measured new learning and visual episodic memory [34]. A number of boxes were shown on the screen. Randomly, two or more of the boxes opened one by one and presented a pattern for 2 s. Patterns were then shown, one at a time, in the screen center, and the participants were instructed to select the box in which the pattern had been displayed.

*RTI.* The RTI test measured divided attention and psychomotor speed [35]. The participants were instructed to hold down a button at the bottom of the screen, followed by quickly tapping inside one of 5 circles at the top of the screen once a yellow stimulus flash appeared. They used the same finger to tap the flashing circle. *SWM.* The SWM task measured manipulation and retention of visuospatial information [36]. The screen showed a number of boxes, where a few contained a hidden token. The participants clicked the empty boxes to locate the tokens and were asked not to go back to a box where one had already been located. Trials increased in difficulty (i.e., number of boxes to search).

### 2.5 Mental health questionnaires

#### 2.5.1 Beck Depression Inventory (BDI)

The BDI assessed symptoms associated with depression over the past two weeks including cognitive (e.g., concentration, pessimism, decisiveness), somatic (e.g., libido, appetite, sleep), and affective symptoms (e.g., sadness, crying, irritability). Each question can be rated in severity from zero to three, where three indicates the highest symptom severity [37]. Adding the scores together provides a total score between 0 and 63. Scores from 0 to 9 indicate minimal depression, 10 to 18 indicate mild depression, 19 to 29 indicate moderate depression, and 30 to 63 indicate severe depression [38]. The BDI has been validated in college students [39].

#### 2.5.2 Beck Anxiety Inventory (BAI)

The BAI assessed anxiety symptoms within the past month. The same assessment scale is used as in the BDI, with four statements in each item, and with a score of three indicating the highest anxiety symptom [40]. The items measure symptoms including physical (e.g., feeling hot, dizziness, difficulty breathing, increased heartrate, sweats), affective (e.g., feelings of fear, nervousness, being scared) and cognitive symptoms (e.g., catastrophizing thoughts). The scores from each item are added together to provide a total score between 0 and 63 [40]. Minimal anxiety is indicated by scores between 0 to 7, scores between 8 to 15 indicate mild anxiety, scores between 16 to 25 indicate moderate anxiety, and scores between 26 to 63 indicate severe anxiety. The BAI has been found to reliably distinguish anxiety from depression in a sample of college students [41].

## 3. Data Analysis

### 3.1 Actigraphy

Actigraphy data were analyzed using ActStudio (version 2.2.0, Condor Instruments, 2022). Only nocturnal sleep was assessed. The sleep parameters estimated were: 1) total sleep time, 2) sleep efficiency (i.e., percentage of time spent asleep of the total time in bed), 3) bedtime, 4) and wake-up time. A minimum of 5 nights of complete actigraphy data were required for each participant to be included in the analyses. The sleep diaries were used to confirm bed and wake-up times. Additionally, we used Clocklab (Version 6.1.11, Actimetrics, Wilmette, USA) to assess circadian parameters of the rest-activity rhythms. Cosinor analysis was used to estimate amplitude, acrophase, intradaily variability (IV), and interdaily stability (IS) of the participants’ daily activity rhythms. Weekdays and weekends were included in the analyses.

### 3.2 PVT

The performance metric calculated for the 5-minute PVT was the slowest 10% reaction times, which is one of the most sensitive PVT measures to sleep disturbance [28,42].

### 3.3 MST

The Memory Recognition scores (REC) for old (i.e., repeat) images were calculated by subtracting the rate of “old” responses provided to foils from the rate of “old” responses provided to repeated images [p(correct old response to lures) - p(false old response to foils)] [43]. The Lure Discrimination Index (LDI) was calculated by subtracting the probability value of “similar” responses provided to the foils from the probability value of “similar” responses provided to the lures [p(correct similar response to lures) - p(false similar response to foils)] [43]. We also assessed performance on L1 and L2 specific lure bins. These lure images are most similar to target images, and thus tax pattern separation the most. For these specific lure bins, we assessed *Accuracy,* as the likelihood judging the lure as similar, and *False Memory Error Rate*, as the likelihood of incorrectly judging the lure as a target image.

### 3.4 CANTAB

The following performance metrics were used in the analyses: *DMS.* We used DMS Pattern Errors (All Delays), which was calculated across all trials with a delay component [44]. The total score was derived from the number of times the participant initially selected the distractor stimulus with different patterns but that had identical color elements. *PAL.* We used the PAL First attempt and PAL Adjusted Errors. PAL First Attempt assessed the number of times a participant chose the correct box on their first attempt while recalling all pattern locations. PAL Adjusted Errors measured the number of times a participant chose the incorrect box, adding an adjustment to allow for a comparison of error rates across all participants [44]. *RTI.* We used the RTI Mean Five-Choice Movement Time, which measured the average time it took for a participant to release the response button and tap the target stimulus [44]. *SWM.* We used SWM Between Errors and SWM Strategy. The SWM Between Errors assessed the number of times a participant re-opened a box where they had previously found a token [44]. The SWM Strategy measured the number of times the participant used the same or different search pattern in trials with 6-8 boxes.

### 3.5 Mental health

We used the total scores of BDI and BAI in the analyses.

### 3.6 Statistical analyses

The relationships between sleep, mental health, and cognition were analyzed using multiple linear regressions in JMP 18.2.1 (JMP Statistical Discovery LLC, North Carolina, USA). Pearson Coefficient analyses assessed relationships before performing the moderation analyses. Data that showed non-normality checked with the Shapiro-Wilks test (p < .05) were run using Spearman’s Rank correlations (ρ). In the regressions, the predictors included mental health, and the moderators included objective sleep measures (Figure 2). Moderation analyses were performed when sleep showed significant associations with the cognitive and/or mental health variables. The relationships between the variables were linear shown by residual plots. The residuals were normally distributed, there was constant variance, and observations were independent shown by the Durbin Watson’s test (*p* > .05). Thus, the assumptions for regression analysis were met. The cut-off used for statistical significance was *p* < 0.05 (two-tailed).

**Figure 2.**
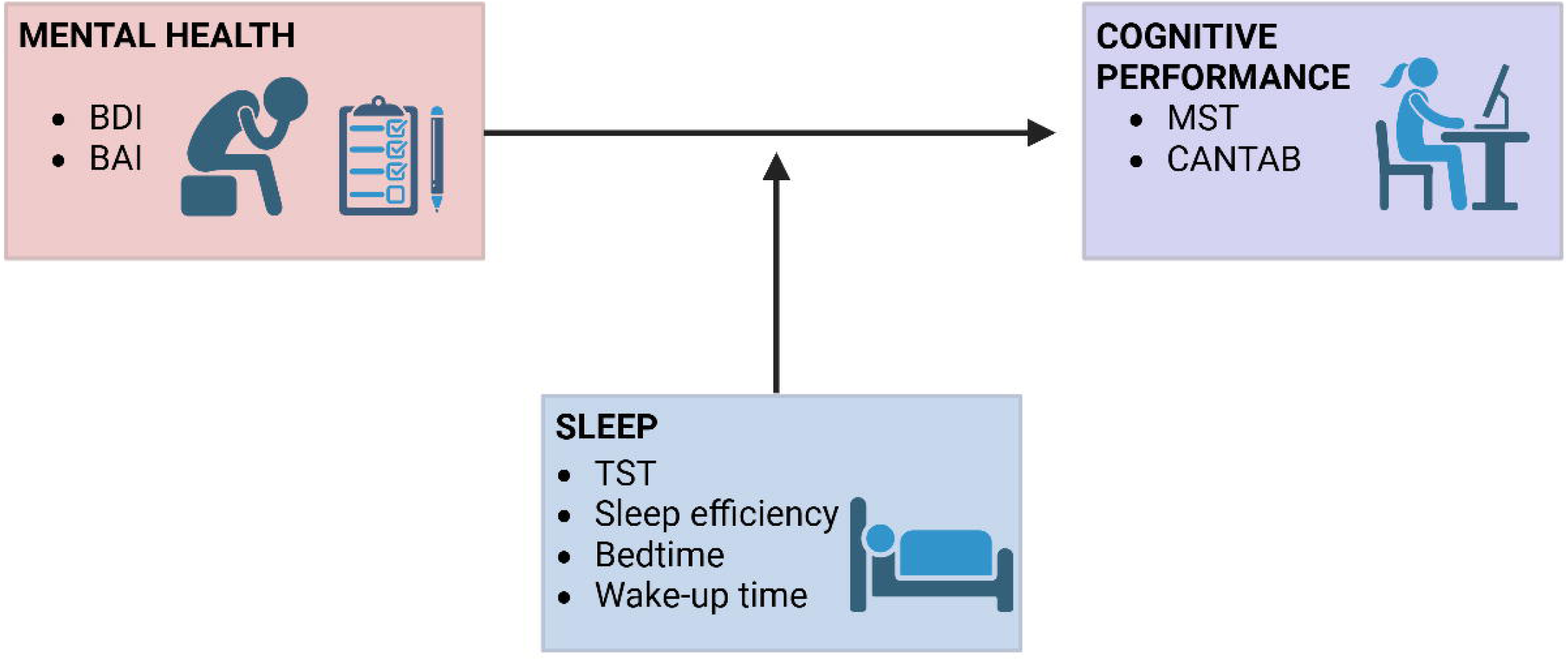
Illustration of the hypothesized moderation relationship between mental health, sleep, and cognitive performance. Abbreviations: BDI, Beck’s Depression Inventory; BAI, Beck’s Anxiety Inventory; TST, total sleep time; MST, Mnemonic Similarity Task; CANTAB, Cambridge Neuropsychological Test Automated Battery.

## 4. Results

### 4.1 Sample characteristics

Of the 89 participants, 9 were excluded from all analyses due to unreliable or missing sleep data (*n* = 5), external distractions during in-person testing session (*n* = 1), malfunction on the CANTAB and incorrect task administration on the MST (*n* = 1), being outside of recruitment age range (*n* = 1), or missing mental health data (*n* = 1). Eighty participants (mean age: 20.13 ± 2.00) were included in the analyses.

For the task-specific analyses, an additional participant was excluded from PVT analyses due to reporting major distractions during task performance. Eleven participants were excluded from MST analyses due to MST malfunction or incorrect test administration (*n* = 5), low effort or not understanding the task indicated by underuse of the “similar” response (*n* = 2) [45], not complying with task instructions (*n* = 2), and a REC score below the cut-off value of .50 (*n* = 2) [32]. Five participants were excluded from CANTAB analyses due to technical error (*n* = 2) and not complying with the protocol (*n* = 3). Twenty-two participants were excluded from Clocklab analyses as they did not have 5 or more days of continuous wrist actigraphy wearing (i.e., removed wrist actigraphy during the day).

### 4.2 Mental health symptoms were common amongst participants

On the BDI, 43.75% of participants scored between 0 and 9, indicating minimal depression, 38.75% scored between 10 and 18, indicating mild depression, 11.25% scored between 19 and 29, indicating moderate depression, and 6.25% scored between 30 and 63, indicating severe depression [38].

On the BAI, 33.75% of participants scored between 0 and 7, indicating minimal anxiety, 26.25% scored between 8 and 15, indicating mild anxiety, 18.75% scored between 16 and 25, indicating moderate anxiety, and 21.25% scored 26 or higher, indicating severe anxiety [40]. The depression and anxiety scores were strongly correlated (ρ(78) *=* .68, *p* < .0001, 95% CI = .52, .79), indicating comorbidity.

### 4.3 Insufficient sleep was common amongst participants

The average total sleep time was 6.69 h (± 0.83) (Table 2), which is less than the recommendations from the American Academy of Sleep Medicine (AASM) and Sleep Research Society (SRS) to avoid the health risks of chronic inadequate sleep [46].

**Table 2.** Summary of sleep and circadian rhythm parameters

|  | Bedtime | Wake-up<br>time | TST (h) | Sleep<br>efficiency (%) | SOL (h) | WASO (h) | Amplitude | Acrophase (h) | IV | IS |
| --- | --- | --- | --- | --- | --- | --- | --- | --- | --- | --- |
| | Mean ± SD | Mean ± SD | Mean ± SD | Mean ± SD | Mean ± SD | Mean ± SD | Mean ± SD | Mean ± SD | Mean ± SD | Mean ± SD |
| **Participants** | 01.51 ± 1.49 | 9.02 ± 1.37 | 6.69 ± 0.83 | 89.22 ± 5.04 | 0.06 ± 0.02 | 0.70 ± 0.39 | 1.63 ± 0.59 | 17.18 ± 1.81 | 1.03 ± 0.29 | 0.39 ± 0.10 |
| Abbreviations: TST, Total Sleep Time; SOL, Sleep Onset Latency; WASO, Wake After Sleep Onset; IV, Intradaily Variability; IS, Interdaily Stability. Bedtime, Wake-up time, TST, SOL, WASO, and acrophase are displayed in decimal time. |  |  |  |  |  |  |  |  |  |  |

### 4.4 Symptoms of depression were associated with later wake-up times

There was a significant correlation between BDI score and wake-up time (ρ(78) *=* .35, *p* = .002, 95% CI = .13, .54) (Figure 3). There were no other significant correlations between the objective sleep measures or circadian parameters and mental health scores (*p >* 0.05). See Supplementary Table S1 for the correlations.

**Figure 3.**
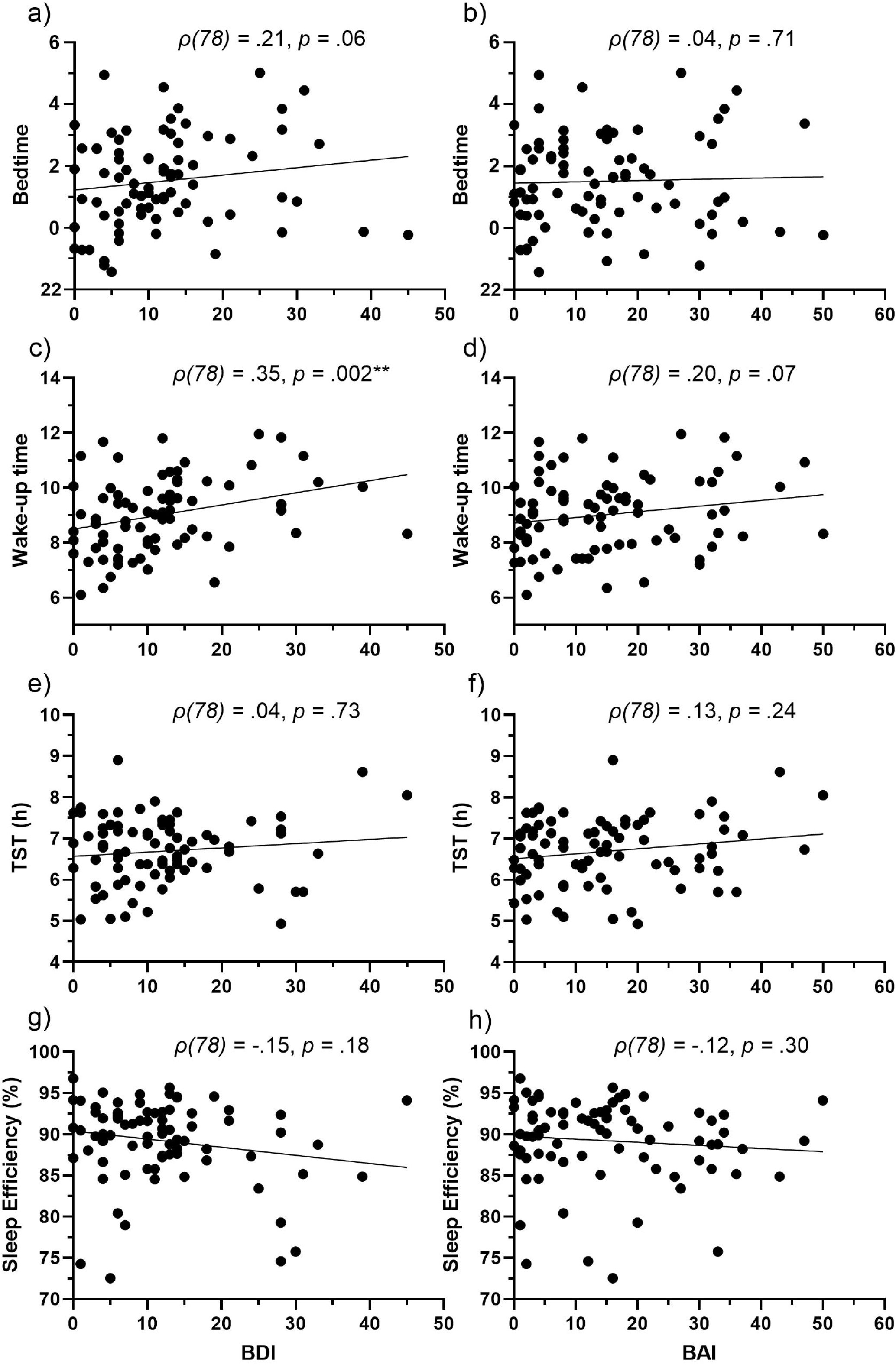
Scatterplots showing the relationship between mental health total scores and sleep: Bedtime (a, b), Wake-up time (c, d), TST (e, f), and sleep efficiency (g, h). Each black dot represents an individual participant. Spearman’s (ρ) correlation, * = *p* < 0.05. Abbreviations: TST, total sleep time; BDI, Beck’s Depression Inventory; BAI, Beck’s Anxiety Inventory.

### 4.5 Symptoms of depression were associated with poorer cognitive performance

There was a significant correlation between BDI total score and PVT 10% slowest reaction time (ρ(77) *=* .26, *p* = .02, 95% CI = .05, .45), such that higher BDI total scores were associated with slower reaction times. There was a significant correlation between BDI total score and CANTAB DMS errors (ρ(71) *=* .24, *p* = .04, 95% CI = .02, .46), such that higher total scores were associated with more errors. No other correlations between BDI and cognitive performance measures were significant (*p >* 0.05) (MST; Figure 4 and 5). See Supplementary Table S1 for the remaining correlations between mental health and cognitive performance.

**Figure 4.**
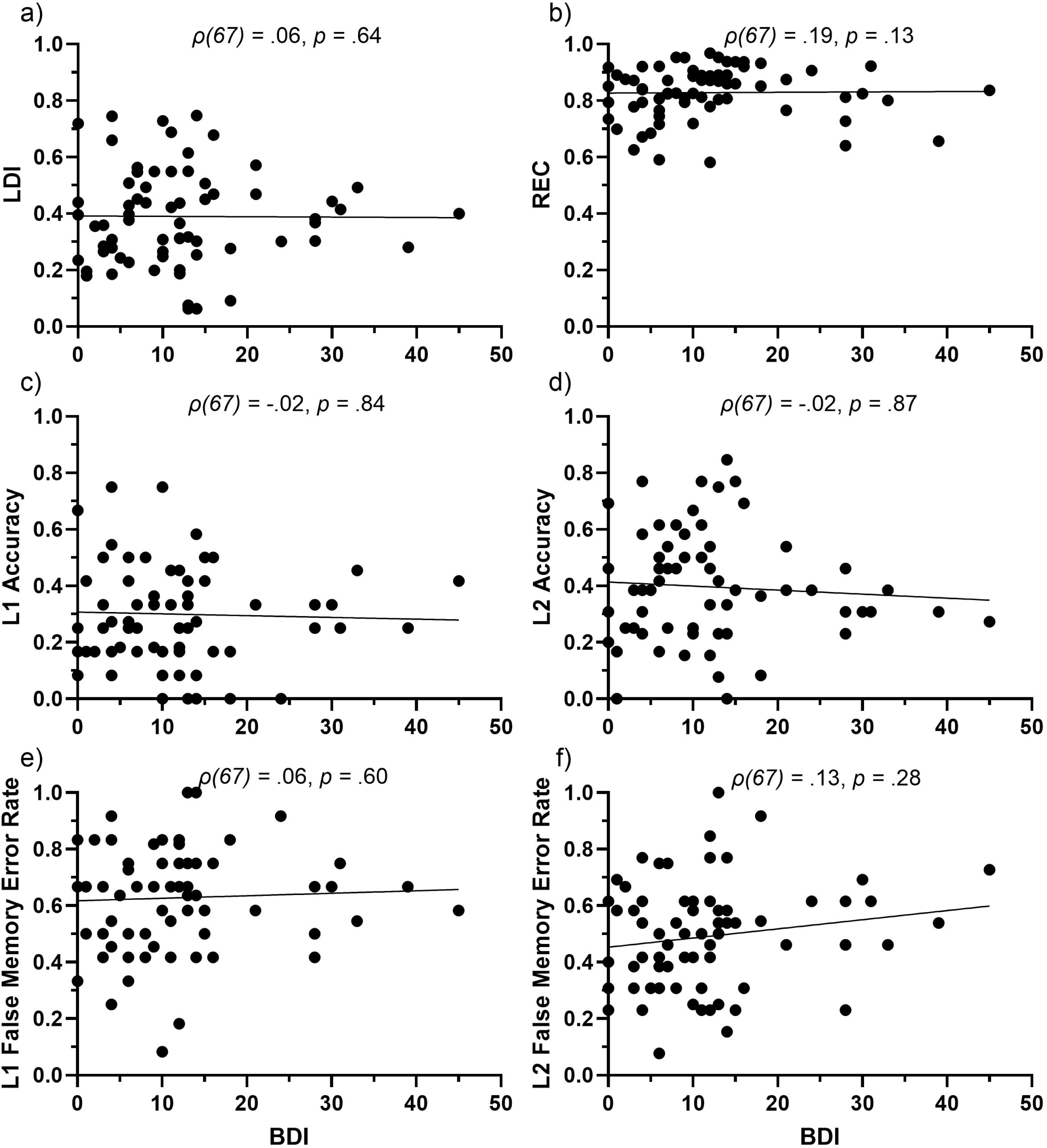
Scatterplots showing the relationship between BDI total score and MST performance: LDI (a), REC (b), L1 Accuracy (c), L2 Accuracy (d), L1 False Memory Error Rate (e), and L2 False Memory Error Rate (f). Each black dot represents an individual participant. Spearman’s (ρ) correlation, * = *p* < 0.05. Abbreviations: BDI, Beck’s Depression Inventory; LDI, Lure Discrimination Index; REC, Recognition Memory; L1, Lure Bin 1; L2, Lure Bin 2.

**Figure 5.**
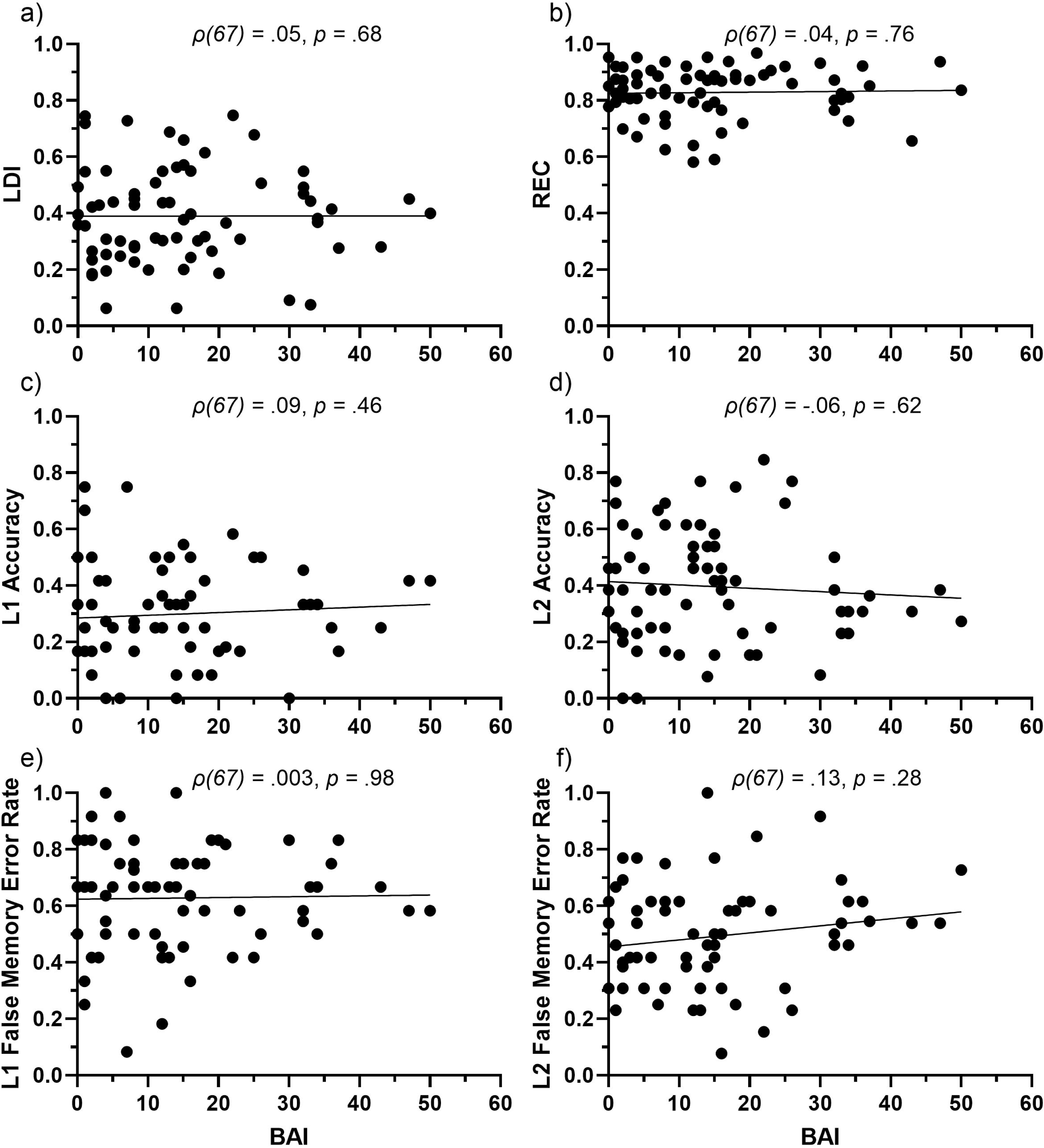
Scatterplots showing the relationship between BAI total score and MST performance: LDI (a), REC (b), L1 Accuracy (c), L2 Accuracy (d), L1 False Memory Error Rate (e), and L2 False Memory Error Rate (f). Each black dot represents an individual participant. Spearman’s (ρ) correlation, * = *p* < 0.05. Abbreviations: BAI, Beck’s Anxiety Inventory; TST, total sleep time; LDI, Lure Discrimination Index; REC, Recognition Memory; L1, Lure Bin 1; L2, Lure Bin 2.

### 4.6 Later bedtime and wake-up time were associated with worse performance on tests of pattern separation

There was a correlation between DMS Pattern Errors (All Delays) and bedtime (ρ(73) *=* .25, *p* = .03, 95% CI = .04, .45) and wake-up time (ρ(73) *=* .28, *p* = .01, 95% CI = .07, .47), such that later bedtimes and wake-up times were associated with more errors. There were also significant correlations between LDI and bedtime (*r*(67) *=* -.29, *p* = .02, 95% CI = -.49, .06) and wake-up time (*r*(67) *=* -.31, *p* = .01, 95% CI = -.51, -.07) (Figure 6). There were significant correlations between bedtime and L2 Accuracy (*r*(67) *=* -.26, *p* = .03, 95% CI = -.47, .02) and L2 False Memory Error Rate (*r*(67) *=* .24, *p* = .047, 95% CI = .003, .45) (Figure 6). See Supplementary Table S2 for the remaining findings. In the circadian analyses, L1 Accuracy was significantly associated with later acrophase (*r*(67) *=* -.29, *p* = .03, 95% CI = -.53, .-01). See Supplementary Table S3 for the remaining results.

**Figure 6.**
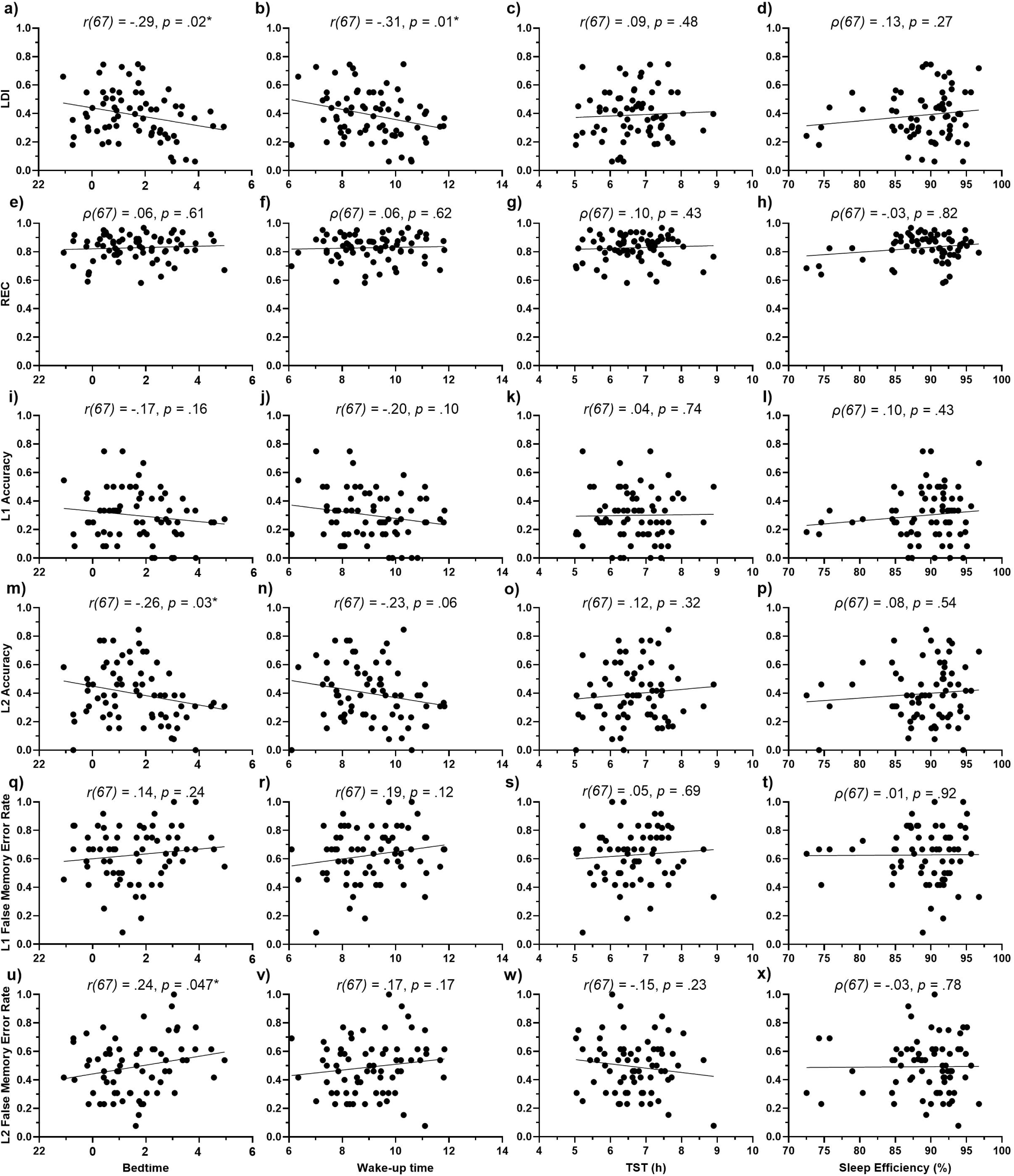
Scatterplots showing the relationship between sleep and MST performance: LDI (a, b, c, d), REC (e, f, g, h), L1 Accuracy (i, j, k, l), L2 Accuracy (m, n, o, p), L1 False Memory Error Rate (q, r, s, t), and L2 False Memory Error Rate (u, v, w, x). Each black dot represents an individual participant. Spearman’s (ρ) correlation, * = *p* < 0.05. Abbreviations: TST, total sleep time; LDI, Lure Discrimination Index; REC, Recognition Memory; L1, Lure Bin 1; L2, Lure Bin 2.

### 4.7 Sleep did not moderate the effect of mental health symptoms on cognitive performance

Sleep did not moderate the relationships between mental health (i.e., BDI and BAI) and cognitive performance (*p >* .05). When analyzing the relationship with BDI, there were direct effects of wake-up time (B = -.04, *p =* .01, 95% CI = -.07, -.01) and bedtime (B = -.03, *p =* .02, 95% CI = -.06, -.01) on LDI. When analyzing the relationship with BAI, there were direct effects of wake-up time on LDI (B = -.04, *p =* .01, 95% CI = -.07, -.01) and DMS (B = .25, *p =* .03, 95% CI = .02, .48), and of bedtime on LDI (B = -.04, *p =* .02, 95% CI = -.06, -.01), and L2 Accuracy (B = -.04, *p =* .04, 95% CI = -.07, .00). See Supplementary Table S4 for the remaining results.

## 5. Discussion

It is well-documented that sleep disturbances, anxiety, and depression can independently and negatively affect cognitive functions [14–17]. In the present study, we investigated whether the sleep disturbances associated with symptoms of anxiety and depression in college students moderates the relationship between mental health and cognitive performance.

Our results showed that more than 50% of the university students who participated in our study reported mild to severe depressive and anxiety symptoms. These findings are consistent with previous research showing that depressive and anxiety symptoms are prevalent in college students [47]. These symptoms of depression and anxiety are often associated with poor sleep [48–50]. In our study, we did not see a relationship between depressive or anxiety symptoms and total sleep time or sleep efficiency, but we did see a relationship between depressive symptoms and the timing of sleep, specifically later bed and wake-up time, which is consistent with previous research [51–55].

We found that later bed and wake-up time were associated with worse cognitive performance, specifically, performance on MST and DMS, which are tasks designed to assess pattern separation. However, total sleep duration and sleep efficiency did not show a relationship with the cognitive measures, and bed and wake-up time did not moderate the relationship between depression and anxiety, and cognitive performance.

The relationships between depression and cognitive performance support previous research [17]. Becker and colleagues (2009) found that participants with a high score on the BDI showed impaired performance on the DMS. Similarly, we found that individuals scoring higher on the BDI showed more errors on the DMS compared to those who had a lower BDI score. We hypothesize that individuals experiencing prolonged stress and/or depressive symptoms may have suppressed hippocampal plasticity, which is important for pattern separation processes [17,20]. We did not find effects of depression on MST performance, which contradicts some previous research [56–59].

Our study has limitations. We did not use a clinical assessment of depression or anxiety but relied on self-report using the BDI and BAI. We used wrist actigraphy and sleep diaries to assess sleep, instead of EEG or polysomnography, thus, we do not have the duration of specific sleep stages or analyses of quantitative aspects of EEG. Our sample included university students who may have different sleep patterns [10] and unique stressors that are different from other young adults, thus, the relationship between sleep, cognition, and mental health that we see in this sample may not generalize beyond the population of college students. Finally, we also included limited control of extraneous variables in this study, thus, effects from medications, caffeine, and different demographic variables may also have impacted the results.

## 6. Conclusion

To summarize, our findings suggest that there may be independent effects of mental health and sleep on cognitive performance in university students. Future research is needed to evaluate whether cognitive tests designed to assess pattern separation are sensitive enough to identify improvements with sleep-targeting interventions or mental health therapeutics.

## Data availability

Data will be available upon request.

## Ethics approval statement

The research study was reviewed and approved by the Research Ethics Board at Simon Fraser University (Protocol #30000539). All participants provided written informed consent to participate in the research. The privacy rights of the participants were observed.

## Acknowledgements

This work was supported by SFU New Faculty Start-up grant (Project Number: 13-2720-00000-N000844, B.A.K.), Canada Research Chairs (Project Number: CRC-2020-00047, B.A.K.), and Canada Foundation for Innovation (CFI-41428, B.A.K.). We would like to thank the following research assistants who assisted with data collection and analyses: Julia Mugliston, Kashish Mehta, Japneet Kaur, Arman Virk, Olivia Braziller, Karthikha Sri Indran, Hannah Chute, Tina Shehata, Vanessa Salzano, Sara Tam, Liliane Tansley, Robert Gibson, and Alex Nash. We would also like to thank Craig Stark (University of California, Irvine) for his invaluable advice and assistance in using the Mnemonic Similarity Task, and Ian Berkovitz (SFU) for helpful statistical consultations.

