## Supplementary Material for "Sleep and mental health independently affect cognitive performance in university students"

| **Table S1.** Correlations between mental health measures and sleep and performance on cognitive tests | | | | | | | | | |
| --- | --- | --- | --- | --- | --- | --- | --- | --- | --- |
|  | **Mental health measures** | | | | | | |  |  |
|  | **BDI** | |  |  |  | **BAI** | |  |  |
|  |  | | **95% CI** | |  |  | | **95% CI** | |
| **Outcomes** | *r/ρ*(df) | *p* | L | U |  | *r/ρ*(df) | *p* | L | U |
| **Sleep**  **(n = 80)** |  |  |  |  |  |  |  |  |  |
| TST | *ρ(78) =* .04 | .73 | -.20 | .28 |  | *ρ(78) =* .13 | .24 | -.10 | .36 |
| SE | *ρ(78) = -*.15 | .18 | -.39 | .09 |  | *ρ(78) = -*.12 | .30 | -.34 | .11 |
| Bedtime | *ρ(78) =* .21 | .06 | -.04 | .44 |  | *ρ(78) =* .04 | .71 | -.20 | .29 |
| Wake-up time | *ρ(78) =* .35 | **.002**** | .13 | .54 |  | *ρ(78) =* .20 | .07 | -.01 | .42 |
| **Circadian parameters**  **(n = 52)** |  |  |  |  |  |  |  |  |  |
| Amplitude | *ρ(56) =* .01 | .94 | -.24 | .26 |  | *ρ(56) =* .13 | .32 | -.14 | .38 |
| Acrophase | *ρ(56) = -*.05 | .72 | -.30 | .19 |  | *ρ(56) = -*.23 | .08 | -.44 | -.00 |
| IV | *ρ(56) = -*.17 | .19 | -.41 | .09 |  | *ρ(56) = -*.19 | .16 | -.44 | .08 |
| IS | *ρ(56) =* .03 | .81 | -.26 | .30 |  | *ρ(56) =* .13 | .33 | -.15 | .40 |
| **PVT**  **(n = 79)** |  |  |  |  |  |  |  |  |  |
| PVT 10% Slowest RT | *ρ(77) =* .26 | **.02*** | .05 | .45 |  | *ρ(77) =* .14 | .22 | -.05 | .33 |
| **MST**  **(n = 69)** |  |  |  |  |  |  |  |  |  |
| LDI | *ρ(67) =* .06 | .64 | -.17 | .29 |  | *ρ(67) =* .05 | .68 | -.18 | .29 |
| REC | *ρ(67) =* .19 | .13 | -.05 | .42 |  | *ρ(67) =* .04 | .76 | -.18 | .27 |
| L1 Accuracy | *ρ(67) =* -.02 | .84 | -.26 | .21 |  | *ρ(67) =* .09 | .46 | -.15 | .34 |
| L2 Accuracy | *ρ(67) =* -.02 | .87 | -.24 | .20 |  | *ρ(67) =* -.06 | .62 | -.28 | .18 |
| L1 False Memory | *ρ(67) =* .06 | .60 | -.18 | .31 |  | *ρ(67) =* .003 | .98 | -.25 | .24 |
| L2 False Memory | *ρ(67) =* .13 | .28 | -.10 | .36 |  | *ρ(67) =* .13 | .28 | -.10 | .37 |
| **CANTAB**  **(n = 75)** |  |  |  |  |  |  |  |  |  |
| DMS Pattern Errors (All Delays) | *ρ(73) =* .24 | **.04*** | .02 | .46 |  | *ρ(73) =* .09 | .43 | -.15 | .31 |
| PAL First Attempt | *ρ(73) =* .11 | .34 | -.12 | .33 |  | *ρ(73) =* .22 | .053 | .01 | .43 |
| PAL Adjusted Errors | *ρ(73) =* -.10 | .39 | -.33 | .12 |  | *ρ(73) =* -.19 | .11 | -.40 | .02 |
| RTI | *ρ(73) =* -.11 | .33 | -.36 | .15 |  | *ρ(73) =* -.06 | .60 | -.31 | .18 |
| SWM Between Errors | *ρ(73) = -*.11 | .35 | -.32 | .13 |  | *ρ(73) = -*.05 | .68 | -.27 | .19 |
| SWM Strategy | *ρ(73) = -*.02 | .86 | -.23 | .19 |  | *ρ(73) = .*03 | .80 | -.19 | .25 |
| Abbreviations: BDI, Beck’s Depression Inventory; BAI, Beck’s Anxiety Inventory; TST, Total Sleep Time; SE, Sleep Efficiency; SOL, Sleep Onset Latency; WASO, Wake After Sleep Onset; IV, Intradaily Variability; IS, Interdaily Stability; PVT, Psychomotor Vigilance Task; RT, Reaction Time; MST, Mnemonic Similarity Task; LDI, Lure Discrimination Index; REC, Recognition Memory; L1, Lure Bin 1; L2, Lure Bin 2; CANTAB, Cambridge Neuropsychological Test Automated Battery; DMS, Delayed Matching to Sample; PAL, Paired Associates Learning; RTI, Reaction Time Inventory; SWM, Spatial Working Memory.  Correlations were run using Pearson’s correlation coefficient (*r*). Correlations with non-normal data were conducted using Spearman’s rank correlation (*ρ*).  * = *p* < .05 (two-tailed). | | | | | | | | | |

| **Table S2.** Correlations between sleep metrics and performance on the cognitive tests | | | | | | | | | | | | | | | | | | | |
| --- | --- | --- | --- | --- | --- | --- | --- | --- | --- | --- | --- | --- | --- | --- | --- | --- | --- | --- | --- |
|  | **Bedtime** | | | |  | **Wake-up time** | | | |  | **TST** | | | |  | **SE** | | | |
|  |  | | **95% CI** | |  |  | | **95% CI** | |  |  | | **95% CI** | |  |  | | **95% CI** | |
| **Cognitive outcomes** | *r/ρ*(df) | *p* | L | U |  | *r/ρ*(df) | *p* | L | U |  | *r/ρ*(df) | *p* | L | U |  | *r/ρ*(df) | *p* | L | U |
| **PVT**  **(n = 79)** |  |  |  |  |  |  |  |  |  |  |  |  |  |  |  |  |  |  |  |
| PVT 10% Slowest RT | *ρ(77) =* .15 | .18 | -.09 | .38 |  | *ρ(77) =* .15 | .18 | -.10 | .38 |  | *ρ(77) = -*.01 | .95 | -.25 | .22 |  | *ρ(77) = -*.20 | .08 | -.39 | .01 |
| **MST**  **(n = 69)** |  |  |  |  |  |  |  |  |  |  |  |  |  |  |  |  |  |  |  |
| LDI | *r(67) =* -.29 | **.02*** | -.49 | .06 |  | *r(67) =* -.31 | **.01*** | -.51 | -.07 |  | *r(67) =* .09 | .48 | -.15 | .32 |  | *ρ(67) =* .13 | .27 | -.10 | .35 |
| REC | *ρ(67) =* .06 | .61 | -.20 | .30 |  | *ρ(67) =* .06 | .62 | -.21 | .31 |  | *ρ(67) =* .10 | .43 | -.16 | .34 |  | *ρ(67) = -*.03 | .82 | -.28 | .20 |
| L1 Accuracy | *r(67) =* -.17 | .16 | -.39 | .07 |  | *r(67) =* -.20 | .10 | -.42 | .04 |  | *r(67) =* .04 | .74 | -.20 | .28 |  | *ρ(67) =* .10 | .43 | -.13 | .32 |
| L2 Accuracy | *r(67) =* -.26 | **.03*** | -.47 | .02 |  | *r(67) =* -.23 | .06 | -.44 | .01 |  | *r(67) =* .12 | .32 | -.12 | .35 |  | *ρ(67) =* .08 | .54 | -.15 | .33 |
| L1 False Memory | *r(67) =* .14 | .24 | -.10 | .37 |  | *r(67) =* .19 | .12 | -.05 | .41 |  | *r(67) =* .05 | .69 | -.19 | .28 |  | *ρ(67) =* .01 | .92 | -.22 | .25 |
| L2 False Memory | *r(67) =* .24 | **.047*** | .003 | .45 |  | *r(67) =* .17 | .17 | -.07 | .39 |  | *r(67) =* -.15 | .23 | -.37 | .09 |  | *ρ(67) =* -.03 | .78 | -.30 | .20 |
| **CANTAB**  **(n = 75)** |  |  |  |  |  |  |  |  |  |  |  |  |  |  |  |  |  |  |  |
| DMS Pattern Errors (All Delays) | *ρ(73) =* .25 | **.03*** | .04 | .45 |  | *ρ(73) =* .28 | **.01*** | .07 | .47 |  | *ρ(73) = -*.01 | .96 | -.22 | .23 |  | *ρ(73) = -*.16 | .16 | -.38 | .06 |
| PAL First Attempt | *ρ(73) =* .03 | .81 | -.21 | .25 |  | *ρ(73) =* -.04 | .73 | -.28 | .20 |  | *ρ(73) = -*.04 | .70 | -.29 | .19 |  | *ρ(73) =* .08 | .51 | -.15 | .30 |
| PAL Adjusted Errors | *ρ(73) =* -.05 | .69 | -.28 | .18 |  | *ρ(73) =* .02 | .84 | -.22 | .26 |  | *ρ(73) =* .05 | .65 | -.17 | .29 |  | *ρ(73) = -*.07 | .54 | -.29 | .15 |
| RTI | *ρ(73) =* -.09 | .44 | -.32 | .14 |  | *ρ(73) =* -.06 | .60 | -.30 | .18 |  | *ρ(73) =* .06 | .64 | -.18 | .30 |  | *ρ(73) =* .08 | .50 | -.15 | .32 |
| SWM Between Errors | *ρ(73) =* -.11 | .36 | -.32 | .10 |  | *ρ(73) =* -.06 | .58 | -.29 | .16 |  | *ρ(73) =* .005 | .97 | -.23 | .26 |  | *ρ(73) = -*.10 | .39 | -.31 | .15 |
| SWM Strategy | *ρ(73) =* .05 | .67 | -.15 | .27 |  | *ρ(73) =* .02 | .83 | -.17 | .24 |  | *ρ(73) =* .01 | .93 | -.21 | .22 |  | *ρ(73) =* .03 | .79 | -.21 | .25 |
| Abbreviations: PVT, Psychomotor Vigilance Task; RT, Reaction Time; MST, Mnemonic Similarity Task; LDI, Lure Discrimination Index; REC, Recognition  Memory; L1, Lure Bin 1; L2, Lure Bin 2; CANTAB, Cambridge Neuropsychological Test Automated Battery; DMS, Delayed Matching to Sample; PAL, Paired  Associates Learning; RTI, Reaction Time Inventory; SWM, Spatial Working Memory.  Correlations were run using Pearson’s correlation coefficient (r). Correlations with non-normal data were conducted using Spearman’s rank correlation (ρ).  * = p < .05 (two-tailed). | | | | | | | | | | | | | | | | | | | |

| **Table S3.** Correlations between performance on cognitive tests and circadian rhythm parameters | | | | | | | | | | | | | | | | | | | | | |
| --- | --- | --- | --- | --- | --- | --- | --- | --- | --- | --- | --- | --- | --- | --- | --- | --- | --- | --- | --- | --- | --- |
|  |  |  | **Circadian rhythm parameters** | | | | | | | | | | | | | | | | | | |
|  | **Amplitude** | | | |  |  |  | **Acrophase** | |  |  |  | **IV** | |  |  |  | **IS** | |  |  |
|  |  | | | | **95% CI** | |  |  | | **95% CI** | |  |  | | **95% CI** | |  |  | | **95% CI** | |
| **Cognitive outcomes** | *r/ρ*(df) | | | *p* | L | U |  | *r/ρ*(df) | *p* | L | U |  | *r/ρ*(df) | *p* | L | U |  | *r/ρ*(df) | *p* | L | U |
| **PVT**  **(n = 57)** |  | | |  |  |  |  |  |  |  |  |  |  |  |  |  |  |  |  |  |  |
| Mean Slowest 10% RT | *ρ*(55) *=* 0.14 | | | .31 | -.14 | .39 |  | *ρ*(55) *=* 0.09 | .49 | -.20 | .37 |  | *ρ*(55) *=* 0.01 | .95 | -.25 | .28 |  | *ρ*(55) *=* 0.16 | .23 | -.11 | .41 |
| **MST**  **(n = 50)** |  | | |  |  |  |  |  |  |  |  |  |  |  |  |  |  |  |  |  |  |
| LDI | *ρ*(48) *=* 0.20 | | | .16 | -.10 | .49 |  | *r*(48) *=* -0.18 | .17 | -.44 | .10 |  | *r*(48) *=* -0.02 | .88 | -.30 | .26 |  | *r*(48) *=* 0.16 | .23 | -.12 | .42 |
| REC | *ρ*(48) = -0.10 | | | .49 | -.38 | .20 |  | *ρ*(48) *=* -0.06 | .67 | -.35 | .25 |  | *ρ*(48) *=* 0.19 | .19 | -.08 | .44 |  | *ρ*(48) *=* -0.16 | .28 | -.41 | .13 |
| L1 Accuracy | *ρ(*48) *=* 0.15 | | | .29 | -.15 | .41 |  | *r*(48) *= -*0.29 | **.03*** | -.53 | -.01 |  | *r*(48) *=* 0.02 | .88 | -.26 | .30 |  | *r*(48) *=* -0.003 | .98 | -.28 | .28 |
| L2 Accuracy | *ρ(*48) *=* 0.21 | | | .14 | -.09 | .48 |  | *r*(48) *=* 0.07 | .59 | -.21 | .34 |  | *r*(48) *= -*0.02 | .90 | -.29 | .26 |  | *r*(48) *=* 0.01 | .94 | -.27 | .29 |
| L1 False Memory Error Rate | *ρ*(48) *= -*0.12 | | | .40 | -.40 | .17 |  | *r*(48) *=* 0.18 | .18 | -.11 | .43 |  | *r*(48) *=* 0.05 | .69 | -.23 | .33 |  | *r*(48) *=* 0.02 | .86 | -.26 | .30 |
| L2 False Memory Error Rate | *ρ*(48) *= -*0.22 | | | .13 | -.47 | .08 |  | *r*(48) *= -*0.04 | .78 | -.31 | .24 |  | *r*(48) *=* 0.001 | .99 | -.28 | .28 |  | *r*(48) *=* 0.01 | .94 | -.27 | .29 |
| **CANTAB**  **(n = 53)** |  | | |  |  |  |  |  |  |  |  |  |  |  |  |  |  |  |  |  |  |
| DMS Pattern Errors (All Delays) | *ρ*(51) *=* 0.06 | | | .69 | -.26 | .35 |  | *ρ*(51) *=* 0.10 | .49 | -.18 | .36 |  | *ρ*(51) *=* 0.13 | .36 | -.15 | .38 |  | *ρ*(51) *= -*0.22 | .12 | -.49 | .09 |
| PAL First Attempt | *ρ*(51) *=* -0.15 | | | .27 | -.42 | .14 |  | *ρ*(51) *=* -0.23 | .10 | -.49 | .09 |  | *ρ*(51) = -0.06 | .67 | -.35 | .23 |  | *ρ*(51) = 0.08 | .55 | -.20 | .34 |
| PAL Adjusted Errors | *ρ*(51) *=* 0.14 | | | .33 | -.15 | .40 |  | *ρ*(51) *=* 0.22 | .12 | -.10 | .49 |  | *ρ*(51) = 0.06 | .65 | -.23 | .35 |  | *ρ*(51) = -0.09 | .51 | -.36 | .18 |
| RTI | *ρ*(51) *=* -0.27 | | | .052 | -.50 | -.01 |  | *ρ*(51) *=* 0.17 | .22 | -.11 | .43 |  | *ρ*(51) = 0.08 | .59 | -.19 | .34 |  | *ρ*(51) = -0.07 | .60 | -.35 | .22 |
| SWM Between Errors | *ρ*(51) *=* 0.08 | | | .56 | -.21 | .34 |  | *ρ*(51) *= -*0*.*18 | .20 | -.45 | .12 |  | *ρ*(51) = 0.12 | .39 | -.13 | .37 |  | *ρ*(51) = 0.08 | .57 | -.20 | .36 |
| SWM Strategy | *ρ*(51) *=* 0.09 | | | .52 | -.19 | .35 |  | *ρ*(51) *=* -0*.*15 | .27 | -.43 | .11 |  | *ρ*(51) = -0.01 | .92 | -.26 | .25 |  | *ρ*(51) = 0.08 | .55 | -.22 | .36 |
| Abbreviations: IV, Intradaily Variability; IS, Interdaily Stability; PVT, Psychomotor Vigilance Task; RT, Reaction Time; MST, Mnemonic Similarity Task; LDI, Lure Discrimination Index; REC, Recognition Memory; L1, Lure Bin 1; L2, Lure Bin 2; DMS, Delayed Matching to Sample; PAL, Paired Associates Learning; RTI, Reaction Time Inventory; SWM, Spatial Working Memory.  Correlations were run using Pearson’s correlation coefficient (*r*). Correlations with non-normal data were conducted using Spearman’s rank correlation (*ρ*).  *Note.* Sample sizes for the circadian analyses are lower than for the sleep analyses because we needed at least 5 consecutive days of wrist actigraphy.  * = *p* < . 05 (two-tailed). | | | | | | | | | | | | | | | | | | | | | |

| **Table S4.** Moderation Analysis: Is the effect of mental health on cognitive performance moderated by sleep? | | | | | | | | | |
| --- | --- | --- | --- | --- | --- | --- | --- | --- | --- |
| **Outcome variables** | **Regressions** | **Df,error** | **B** | **Std error** | **t ratio** | ***p*** | **Lower 95%** | **Upper 95%** | **VIF** |
| **BDI & Wake-up time** |  |  |  |  |  |  |  |  |  |
| **LDI** |  |  |  |  |  |  |  |  |  |
|  | Intercept |  | 1.00 | 0.27 | 3.66 | .001* | 0.45 | 1.54 | . |
|  | Age | 1,63 | -0.01 | 0.01 | -1.10 | .27 | -0.03 | 0.01 | 1.15 |
|  | Sex | 1,63 | 0.02 | 0.02 | 0.77 | .44 | -0.03 | 0.06 | 1.25 |
|  | BDI | 1,63 | 0.00 | 0.00 | 0.03 | .97 | 0.00 | 0.00 | 1.26 |
|  | Wake-up time | 1,63 | -0.04 | 0.02 | -2.60 | **.01*** | -0.07 | -0.01 | 1.14 |
|  | BDI*Wake-up time | 1,63 | 0.00 | 0.00 | 0.23 | .82 | 0.00 | 0.00 | 1.08 |
| **L1 Accuracy** |  |  |  |  |  |  |  |  |  |
|  | Intercept |  | 0.50 | 0.29 | 1.73 | .09 | -0.08 | 1.09 | . |
|  | Age | 1,63 | 0.00 | 0.01 | 0.11 | .92 | -0.02 | 0.02 | 1.15 |
|  | Sex | 1,63 | 0.00 | 0.02 | -0.11 | .91 | -0.05 | 0.05 | 1.25 |
|  | BDI | 1,63 | 0.00 | 0.00 | 0.26 | .80 | 0.00 | 0.01 | 1.26 |
|  | Wake-up time | 1,63 | -0.03 | 0.02 | -1.48 | .14 | -0.06 | 0.01 | 1.14 |
|  | BDI*Wake-up time | 1,63 | 0.00 | 0.00 | -0.57 | .57 | 0.00 | 0.00 | 1.08 |
| **L2 Accuracy** |  |  |  |  |  |  |  |  |  |
|  | Intercept |  | 0.89 | 0.33 | 2.69 | .01* | 0.23 | 1.55 | . |
|  | Age | 1,63 | -0.01 | 0.01 | -0.62 | .54 | -0.03 | 0.02 | 1.15 |
|  | Sex | 1,63 | 0.02 | 0.03 | 0.80 | .43 | -0.03 | 0.08 | 1.25 |
|  | BDI | 1,63 | 0.00 | 0.00 | -0.44 | .66 | -0.01 | 0.00 | 1.26 |
|  | Wake-up time | 1,63 | -0.04 | 0.02 | -1.81 | .08 | -0.07 | 0.00 | 1.14 |
|  | BDI*Wake-up time | 1,63 | 0.00 | 0.00 | -0.26 | .80 | 0.00 | 0.00 | 1.08 |
| **L1 False Memory Error Rate** |  |  |  |  |  |  |  |  |  |
|  | Intercept |  | 0.46 | 0.31 | 1.48 | .14 | -0.16 | 1.09 | . |
|  | Age | 1,63 | 0.00 | 0.01 | -0.35 | .73 | -0.03 | 0.02 | 1.15 |
|  | Sex | 1,63 | 0.01 | 0.03 | 0.29 | .77 | -0.04 | 0.06 | 1.25 |
|  | BDI | 1,63 | 0.00 | 0.00 | -0.33 | .74 | -0.01 | 0.00 | 1.26 |
|  | Wake-up time | 1,63 | 0.03 | 0.02 | 1.51 | .14 | -0.01 | 0.06 | 1.14 |
|  | BDI*Wake-up time | 1,63 | 0.00 | 0.00 | 1.03 | .31 | 0.00 | 0.01 | 1.08 |
| **L2 False Memory Error Rate** |  |  |  |  |  |  |  |  |  |
|  | Intercept |  | 0.37 | 0.32 | 1.15 | .25 | -0.27 | 1.00 | . |
|  | Age | 1,63 | 0.00 | 0.01 | -0.26 | .79 | -0.03 | 0.02 | 1.15 |
|  | Sex | 1,63 | -0.02 | 0.03 | -0.77 | .44 | -0.07 | 0.03 | 1.25 |
|  | BDI | 1,63 | 0.00 | 0.00 | 1.10 | .27 | 0.00 | 0.01 | 1.26 |
|  | Wake-up time | 1,63 | 0.02 | 0.02 | 0.93 | .35 | -0.02 | 0.06 | 1.14 |
|  | BDI*Wake-up time | 1,63 | 0.00 | 0.00 | 0.11 | .92 | 0.00 | 0.00 | 1.08 |
| **DMS Pattern Errors (All Delays)** |  |  |  |  |  |  |  |  |  |
|  | Intercept |  | 1.74 | 1.97 | 0.88 | .38 | -2.19 | 5.67 | . |
|  | Age | 1,69 | -0.13 | 0.08 | -1.60 | .11 | -0.30 | 0.03 | 1.18 |
|  | Sex | 1,69 | -0.16 | 0.18 | -0.88 | .38 | -0.51 | 0.20 | 1.32 |
|  | BDI | 1,69 | 0.03 | 0.02 | 1.63 | .11 | -0.01 | 0.07 | 1.25 |
|  | Wake-up time | 1,69 | 0.21 | 0.12 | 1.83 | .07 | -0.02 | 0.44 | 1.14 |
|  | BDI*Wake-up time | 1,69 | -0.02 | 0.01 | -1.38 | .17 | -0.04 | 0.01 | 1.08 |
| **BDI & Bedtime** |  |  |  |  |  |  |  |  |  |
| **LDI** |  |  |  |  |  |  |  |  |  |
|  | Intercept |  | 1.52 | 0.46 | 3.31 | .002* | 0.60 | 2.44 | . |
|  | Age | 1,63 | -0.01 | 0.01 | -1.11 | .27 | -0.03 | 0.01 | 1.16 |
|  | Sex | 1,63 | 0.01 | 0.02 | 0.46 | .65 | -0.03 | 0.06 | 1.19 |
|  | BDI | 1,63 | 0.00 | 0.00 | -0.24 | .81 | -0.01 | 0.00 | 1.23 |
|  | Bedtime | 1,63 | -0.03 | 0.01 | -2.37 | **.02*** | -0.06 | -0.01 | 1.02 |
|  | BDI*Bedtime | 1,63 | 0.00 | 0.00 | 0.72 | .48 | 0.00 | 0.00 | 1.07 |
| **L1 Accuracy** |  |  |  |  |  |  |  |  |  |
|  | Intercept |  | 0.74 | 0.49 | 1.50 | .14 | -0.25 | 1.72 | . |
|  | Age | 1,63 | 0.00 | 0.01 | 0.07 | .95 | -0.02 | 0.02 | 1.16 |
|  | Sex | 1,63 | 0.00 | 0.02 | -0.20 | .84 | -0.05 | 0.04 | 1.19 |
|  | BDI | 1,63 | 0.00 | 0.00 | -0.21 | .84 | -0.01 | 0.00 | 1.23 |
|  | Bedtime | 1,63 | -0.02 | 0.02 | -1.11 | .27 | -0.05 | 0.01 | 1.02 |
|  | BDI*Bedtime | 1,63 | 0.00 | 0.00 | -0.38 | .71 | 0.00 | 0.00 | 1.07 |
| **L2 Accuracy** |  |  |  |  |  |  |  |  |  |
|  | Intercept |  | 1.51 | 0.55 | 2.75 | .01* | 0.41 | 2.61 | . |
|  | Age | 1,63 | -0.01 | 0.01 | -0.72 | .48 | -0.03 | 0.02 | 1.16 |
|  | Sex | 1,63 | 0.02 | 0.03 | 0.67 | .51 | -0.04 | 0.07 | 1.19 |
|  | BDI | 1,63 | 0.00 | 0.00 | -0.83 | .41 | -0.01 | 0.00 | 1.23 |
|  | Bedtime | 1,63 | -0.04 | 0.02 | -2.02 | **.048*** | -0.07 | 0.00 | 1.02 |
|  | BDI*Bedtime | 1,63 | 0.00 | 0.00 | 0.01 | .99 | 0.00 | 0.00 | 1.07 |
| **L1 False Memory Error Rate** |  |  |  |  |  |  |  |  |  |
|  | Intercept |  | 0.29 | 0.53 | 0.54 | .59 | -0.76 | 1.34 | . |
|  | Age | 1,63 | 0.00 | 0.01 | -0.28 | .78 | -0.03 | 0.02 | 1.16 |
|  | Sex | 1,63 | 0.01 | 0.03 | 0.35 | .73 | -0.04 | 0.06 | 1.19 |
|  | BDI | 1,63 | 0.00 | 0.00 | 0.41 | .68 | 0.00 | 0.01 | 1.23 |
|  | Bedtime | 1,63 | 0.02 | 0.02 | 0.92 | .36 | -0.02 | 0.05 | 1.02 |
|  | BDI*Bedtime | 1,63 | 0.00 | 0.00 | 1.22 | .23 | 0.00 | 0.00 | 1.07 |
| **L2 False Memory Error Rate** |  |  |  |  |  |  |  |  |  |
|  | Intercept |  | -0.30 | 0.52 | -0.57 | .57 | -1.34 | 0.74 | . |
|  | Age | 1,63 | 0.00 | 0.01 | -0.11 | .91 | -0.03 | 0.02 | 1.16 |
|  | Sex | 1,63 | -0.02 | 0.03 | -0.73 | .47 | -0.07 | 0.03 | 1.19 |
|  | BDI | 1,63 | 0.00 | 0.00 | 1.38 | .17 | 0.00 | 0.01 | 1.23 |
|  | Bedtime | 1,63 | 0.03 | 0.02 | 1.83 | .07 | 0.00 | 0.06 | 1.02 |
|  | BDI*Bedtime | 1,63 | 0.00 | 0.00 | 0.19 | .85 | 0.00 | 0.00 | 1.07 |
| **DMS Pattern Errors (All Delays)** |  |  |  |  |  |  |  |  |  |
|  | Intercept |  | -1.12 | 3.16 | -0.36 | .72 | -7.42 | 5.18 | . |
|  | Age | 1,69 | -0.14 | 0.08 | -1.62 | .11 | -0.30 | 0.03 | 1.17 |
|  | Sex | 1,69 | -0.09 | 0.18 | -0.53 | .60 | -0.45 | 0.26 | 1.27 |
|  | BDI | 1,69 | 0.03 | 0.02 | 1.54 | .13 | -0.01 | 0.06 | 1.13 |
|  | Bedtime | 1,69 | 0.19 | 0.11 | 1.79 | .08 | -0.02 | 0.40 | 1.04 |
|  | BDI*Bedtime | 1,69 | -0.01 | 0.01 | -0.84 | .40 | -0.03 | 0.01 | 1.04 |
| **BAI & Wake-up time** |  |  |  |  |  |  |  |  |  |
| **LDI** |  |  |  |  |  |  |  |  |  |
|  | Intercept |  | 1.00 | 0.27 | 3.64 | .001* | 0.45 | 1.54 | . |
|  | Age | 1,63 | -0.01 | 0.01 | -1.09 | .28 | -0.03 | 0.01 | 1.16 |
|  | Sex | 1,63 | 0.02 | 0.02 | 0.86 | .39 | -0.03 | 0.06 | 1.23 |
|  | BAI | 1,63 | 0.00 | 0.00 | -0.04 | .97 | 0.00 | 0.00 | 1.18 |
|  | Wake-up time | 1,63 | -0.04 | 0.02 | -2.65 | **.01*** | -0.07 | -0.01 | 1.10 |
|  | BAI*Wake-up time | 1,63 | 0.00 | 0.00 | 0.51 | .61 | 0.00 | 0.00 | 1.09 |
| **L1 Accuracy** |  |  |  |  |  |  |  |  |  |
|  | Intercept |  | 0.44 | 0.29 | 1.51 | .14 | -0.14 | 1.02 | . |
|  | Age | 1,63 | 0.00 | 0.01 | 0.28 | .78 | -0.02 | 0.03 | 1.16 |
|  | Sex | 1,63 | 0.00 | 0.02 | 0.08 | .94 | -0.05 | 0.05 | 1.23 |
|  | BAI | 1,63 | 0.00 | 0.00 | 0.75 | .46 | 0.00 | 0.01 | 1.18 |
|  | Wake-up time | 1,63 | -0.03 | 0.02 | -1.49 | .14 | -0.06 | 0.01 | 1.10 |
|  | BAI*Wake-up time | 1,63 | 0.00 | 0.00 | 0.89 | .38 | 0.00 | 0.00 | 1.09 |
| **L2 Accuracy** |  |  |  |  |  |  |  |  |  |
|  | Intercept |  | 0.89 | 0.33 | 2.70 | .01* | 0.23 | 1.56 | . |
|  | Age | 1,63 | -0.01 | 0.01 | -0.65 | .52 | -0.03 | 0.02 | 1.16 |
|  | Sex | 1,63 | 0.02 | 0.03 | 0.84 | .41 | -0.03 | 0.08 | 1.23 |
|  | BAI | 1,63 | 0.00 | 0.00 | -0.59 | .55 | -0.01 | 0.00 | 1.18 |
|  | Wake-up time | 1,63 | -0.03 | 0.02 | -1.81 | .07 | -0.07 | 0.00 | 1.10 |
|  | BAI*Wake-up time | 1,63 | 0.00 | 0.00 | 0.05 | .96 | 0.00 | 0.00 | 1.09 |
| **L1 False Memory Error Rate** |  |  |  |  |  |  |  |  |  |
|  | Intercept |  | 0.49 | 0.32 | 1.56 | .12 | -0.14 | 1.13 | . |
|  | Age | 1,63 | 0.00 | 0.01 | -0.35 | .73 | -0.03 | 0.02 | 1.16 |
|  | Sex | 1,63 | 0.00 | 0.03 | 0.03 | .98 | -0.05 | 0.05 | 1.23 |
|  | BAI | 1,63 | 0.00 | 0.00 | -0.12 | .90 | 0.00 | 0.00 | 1.18 |
|  | Wake-up time | 1,63 | 0.02 | 0.02 | 1.37 | .18 | -0.01 | 0.06 | 1.10 |
|  | BAI*Wake-up time | 1,63 | 0.00 | 0.00 | -0.38 | .71 | 0.00 | 0.00 | 1.09 |
| **L2 False Memory Error Rate** |  |  |  |  |  |  |  |  |  |
|  | Intercept |  | 0.36 | 0.32 | 1.12 | .27 | -0.28 | 0.99 | . |
|  | Age | 1,63 | 0.00 | 0.01 | -0.23 | .82 | -0.03 | 0.02 | 1.16 |
|  | Sex | 1,63 | -0.02 | 0.03 | -0.77 | .45 | -0.07 | 0.03 | 1.23 |
|  | BAI | 1,63 | 0.00 | 0.00 | 1.21 | .23 | 0.00 | 0.01 | 1.18 |
|  | Wake-up time | 1,63 | 0.02 | 0.02 | 0.97 | .33 | -0.02 | 0.05 | 1.10 |
|  | BAI*Wake-up time | 1,63 | 0.00 | 0.00 | -0.32 | .75 | 0.00 | 0.00 | 1.09 |
| **DMS Pattern Errors (All Delays)** |  |  |  |  |  |  |  |  |  |
|  | Intercept |  | 1.51 | 2.02 | 0.75 | .46 | -2.53 | 5.55 | . |
|  | Age | 1,69 | -0.13 | 0.08 | -1.58 | .12 | -0.30 | 0.04 | 1.19 |
|  | Sex | 1,69 | -0.13 | 0.19 | -0.69 | .49 | -0.50 | 0.24 | 1.42 |
|  | BAI | 1,69 | 0.01 | 0.01 | 0.87 | .39 | -0.02 | 0.04 | 1.20 |
|  | Wake-up time | 1,69 | 0.25 | 0.11 | 2.21 | **.03*** | 0.02 | 0.48 | 1.08 |
|  | BAI*Wake-up time | 1,69 | 0.00 | 0.01 | -0.52 | .60 | -0.02 | 0.01 | 1.11 |
| **BAI & Bedtime** |  |  |  |  |  |  |  |  |  |
| **LDI** |  |  |  |  |  |  |  |  |  |
|  | Intercept |  | 1.54 | 0.47 | 3.31 | .002* | 0.61 | 2.48 | . |
|  | Age | 1,63 | -0.01 | 0.01 | -1.14 | .26 | -0.03 | 0.01 | 1.17 |
|  | Sex | 1,63 | 0.01 | 0.02 | 0.45 | .65 | -0.03 | 0.05 | 1.16 |
|  | BAI | 1,63 | 0.00 | 0.00 | -0.39 | .70 | 0.00 | 0.00 | 1.12 |
|  | Bedtime | 1,63 | -0.04 | 0.01 | -2.37 | **.02*** | -0.06 | -0.01 | 1.03 |
|  | BAI*Bedtime | 1,63 | 0.00 | 0.00 | 0.45 | .66 | 0.00 | 0.00 | 1.04 |
| **L1 Accuracy** |  |  |  |  |  |  |  |  |  |
|  | Intercept |  | 0.67 | 0.50 | 1.35 | .18 | -0.32 | 1.66 | . |
|  | Age | 1,63 | 0.00 | 0.01 | 0.25 | .80 | -0.02 | 0.03 | 1.17 |
|  | Sex | 1,63 | -0.01 | 0.02 | -0.33 | .74 | -0.06 | 0.04 | 1.16 |
|  | BAI | 1,63 | 0.00 | 0.00 | 0.69 | .49 | 0.00 | 0.00 | 1.12 |
|  | Bedtime | 1,63 | -0.02 | 0.02 | -1.10 | .27 | -0.05 | 0.01 | 1.03 |
|  | BAI*Bedtime | 1,63 | 0.00 | 0.00 | 0.07 | .94 | 0.00 | 0.00 | 1.04 |
| **L2 Accuracy** |  |  |  |  |  |  |  |  |  |
|  | Intercept |  | 1.55 | 0.55 | 2.80 | .01* | 0.44 | 2.66 | . |
|  | Age | 1,63 | -0.01 | 0.01 | -0.76 | .45 | -0.04 | 0.02 | 1.17 |
|  | Sex | 1,63 | 0.02 | 0.03 | 0.66 | .51 | -0.04 | 0.07 | 1.16 |
|  | BAI | 1,63 | 0.00 | 0.00 | -0.94 | .35 | -0.01 | 0.00 | 1.12 |
|  | Bedtime | 1,63 | -0.04 | 0.02 | -2.08 | **.04*** | -0.07 | 0.00 | 1.03 |
|  | BAI*Bedtime | 1,63 | 0.00 | 0.00 | 0.30 | .76 | 0.00 | 0.00 | 1.04 |
| **L1 False Memory Error Rate** |  |  |  |  |  |  |  |  |  |
|  | Intercept |  | 0.30 | 0.54 | 0.57 | .57 | -0.77 | 1.38 | . |
|  | Age | 1,63 | 0.00 | 0.01 | -0.35 | .73 | -0.03 | 0.02 | 1.17 |
|  | Sex | 1,63 | 0.01 | 0.03 | 0.38 | .71 | -0.04 | 0.06 | 1.16 |
|  | BAI | 1,63 | 0.00 | 0.00 | 0.03 | .97 | 0.00 | 0.00 | 1.12 |
|  | Bedtime | 1,63 | 0.02 | 0.02 | 0.94 | .35 | -0.02 | 0.05 | 1.03 |
|  | BAI*Bedtime | 1,63 | 0.00 | 0.00 | 0.38 | .70 | 0.00 | 0.00 | 1.04 |
| **L2 False Memory Error Rate** |  |  |  |  |  |  |  |  |  |
|  | Intercept |  | -0.36 | 0.53 | -0.68 | .50 | -1.41 | 0.69 | . |
|  | Age | 1,63 | 0.00 | 0.01 | -0.06 | .95 | -0.03 | 0.02 | 1.17 |
|  | Sex | 1,63 | -0.02 | 0.03 | -0.70 | .48 | -0.07 | 0.03 | 1.16 |
|  | BAI | 1,63 | 0.00 | 0.00 | 1.41 | .16 | 0.00 | 0.01 | 1.12 |
|  | Bedtime | 1,63 | 0.03 | 0.02 | 1.94 | .06 | 0.00 | 0.07 | 1.03 |
|  | BAI*Bedtime | 1,63 | 0.00 | 0.00 | -0.51 | .61 | 0.00 | 0.00 | 1.04 |
| **DMS Pattern Errors (All Delays)** |  |  |  |  |  |  |  |  |  |
|  | Intercept |  | -1.38 | 3.24 | -0.42 | .67 | -7.84 | 5.09 | . |
|  | Age | 1,69 | -0.13 | 0.09 | -1.56 | .12 | -0.30 | 0.04 | 1.19 |
|  | Sex | 1,69 | -0.07 | 0.18 | -0.35 | .72 | -0.43 | 0.30 | 1.35 |
|  | BAI | 1,69 | 0.01 | 0.01 | 1.05 | .30 | -0.01 | 0.04 | 1.14 |
|  | Bedtime | 1,69 | 0.20 | 0.11 | 1.87 | .07 | -0.01 | 0.41 | 1.04 |
|  | BAI*Bedtime | 1,69 | 0.00 | 0.01 | -0.12 | .90 | -0.02 | 0.01 | 1.09 |

Abbreviations: BDI, Beck’s Depression Inventory; BAI, Beck’s Anxiety Inventory; LDI, Lure Discrimination Index; L2, Lure Bin 2; DMS, Delayed Matching to Sample; VIF, Variation Inflation Factor.

* = *p* < .05 (two-tailed).
